# Exosome-Mediated mRNA Delivery For SARS-CoV-2 Vaccination

**DOI:** 10.1101/2020.11.06.371419

**Authors:** Shang Jui Tsai, Chenxu Guo, Alanna Sedgwick, Saravana Kanagavelu, Justin Nice, Sanjana Shetty, Connie Landaverde, Nadia A. Atai, Stephen J. Gould

## Abstract

Expression-dependent, Spike-only vaccines have been developed, deployed, and shown to be effective in the fight against SARS-CoV-2. However, additional approaches to vaccine development may be needed to meet existing and future challenges posed by emerging Spike variant strains, as well as a likely need for different antigen-delivery systems that are safe and effective for regular, periodic re-administration. We report here the development of mRNA-loaded exosomes, demonstrate that they can mediate the functional expression of heterologous proteins *in vitro* and *in vivo*, and have fewer adverse effects than comparable doses of lipid nanoparticles. Furthermore, we applied this approach to the development of an exosome-based, multiplexed mRNA vaccine that drives expression of immunogenic SARS-CoV-2 Nucleocapsid and Spike proteins. This vaccine elicited long-lasting cellular and humoral responses to Nucleocapsid and to Spike, demonstrating that exosome-based mRNA formulations represent a previously unexplored platform in the fight against COVID-19 and other infectious diseases.

## Introduction

Severe acute respiratory syndrome coronavirus 2 (SARS-CoV-2) is the causative agent of COVID-19 ^1, 2^. COVID-19 typically presents with symptoms common to many respiratory infections, including fever and cough, but in many cases progresses to more severe disease that may include acute respiratory distress, disseminated disease, and death ^34–8^. SARS-CoV-2 entered the human population in late 2019 as the result of a zoonotic leap and is most closely related to coronaviruses endemic to bats (Chiroptera)^9^. SARS-CoV-2 is the third recent zoonotic betacoronavirus to enter the human population, the others being responsible for the outbreaks of severe acute respiratory syndrome (SARS-CoV) in 2002 ^10^ and middle east respiratory syndrome (MERS-CoV) in 2012 (Memish et al., 2013), indicative of a generally susceptibility of human populations to coronavirus zoonoses. These zoonoses are more distantly to human endemic betacoronaviruses (OC43, HKU1, etc.) that also cause respiratory infections of milder effect ^11^). While SARS-CoV-2 infection is associated with lower mortality than SARS-CoV or MERS-CoV, SARS-CoV-2 initially displayed a higher rate of transmission, quickly became a major cause of morbidity and mortality worldwide (https://www.cdc.gov/coronavirus/2019-ncov/hcp/clinical-guidance-management-patients.html)(coronavirus.jhu.edu), and has continued to evolve into numerous variant strains that display even more-elevated rates of transmission and an emerging resistance to antibody-based neutralization ^12–15^.

SARS-CoV-2 enters host cells via a multistep pathway that begins with binding between the Spike protein on the virus surface and its cognate receptor proteins on the host cell surface. These include angiotensin-converting enzyme II (ACE2) ^1, 16, 17^, neuropilin-1 ^18, 19^, and perhaps also CD147 ^20^. Following virus-cell binding, host cell proteases (e.g. TMPRSS2 ^16^, cathepsins ^21^, etc.) cleave Spike, potentiating Spike-catalyzed fusion between the viral and cellular membranes and functional infection of the host cell. Not surprisingly, SARS-CoV-2 receptors and proteases are expressed within the respiratory tract, consistent with its respiratory mode of transmission ^22^. However, they are also expressed in many other cell types, allowing SARS-CoV-2 to spread within the body and impact multiple organ systems (brain, heart, gastrointestinal tract, circulatory system, immune system, etc. ^18, 19, 23–26^).

Following virus-cell membrane fusion, the viral genomic RNA (gRNA) is translated to generate two large polyproteins, open reading frame 1 (orf1a) and orf1ab, which are processed to release 16 nonstructural proteins (nsp1-16) ^27^. These early proteins prime the host cell for virus replication and mediate the synthesis of subgenomic viral RNAs and their unique protein products. These include a dozen or more additional proteins, including the SARS-CoV-2 structural proteins Nucleocapsid (N), Spike (S), Membrane (M), and Envelope (E). Spike, Membrane and Envelope are integral membrane proteins, co-translationally translocated into the endoplasmic reticulum (ER), that subsequently drive virion formation while also incorporating the Nucleocapsid and its bound gRNA as well as some other ancillary proteins ^28, 29^, with virus release via lysosomal exocytosis (Ghosh et al., 2020) []. The released viral particles are ∼100 nm diameter, display prominent Spike protrusions from the cell surface and a lumen containing Nucleocapsid-gRNA complexes^30^. SARS-CoV-2 biogenesis also involves extensive processing of its Spike protein at a polybasic site, generating S1 and S2 forms of Spike, with the N-terminal, receptor-binding S1 fragment bound non-covalently to the fusogenic, membrane-anchored S2 fragment ^16, 31, 32^.

Vaccine design should mirror, and ideally improve on, the correlates of protective immunity that arise from natural infections. It is now well-established that SARS-CoV-2 infection generates potent cellular and humoral immune responses to viral proteins that in most cases reverse the course of disease, clear the viral infection, and confer resistance to reinfection in both people and in animal models ^33–36, 37–39^. Disease-preventing vaccines have previously been developed for animal coronaviruses ^40^ and have been successfully developed and deployed for SARS-CoV-2 ^41–47^, a development that is likely to save millions of lives. However, these first-generation SARS-CoV-2 vaccines only elicit immunity to a single viral protein, Spike, the rapid evolution of which may impair vaccine efficacy ^12–15^. Furthermore, the Spike-only vaccine approach ignores the fact that a primary correlate of immunity in COVID-19 patients is the array of potent immune reactions to the SARS-CoV-2 Nucleocapsid protein ^48^.

Here we describe an expression-dependent SARS-CoV-2 vaccine that combines exosome-based delivery, multiplexed mRNA formulation, induction of immunity to both Spike and Nucleocapsid, and antigen design that involves expressing Nucleocapsid in a form designed for improved antigen presentation. Exosomes are small extracellular vesicles (sEVs) of ∼30-150 nm in diameter that are made by all cells, abundant in all biofluids, and mediate intercellular transmission of signals and macromolecules, including RNAs ^49^. Allogenic exosome transplantations and transfusions have been practices in one form or another for more than a century and have never been associated with any adverse effects. Moreover, exosomes have already been shown effective for delivery of RNA-based therapeutics ^50, 51^. The remainder of this report describes the production of engineered exosome/mRNA formulations, their ability to drive protein expression in cultured cells and animals, their improved safety relative to LNPs, and their use as a multiplexed, exosome-based SARS-CoV-2 vaccine that elicited immunity to multiple viral antigens, including Nucleocapsid as well as Spike.

## Results

### Exosomes display robust ability to deliver functional mRNAs in vitro and in vivo

Exosomes are capable of delivering functional RNAs to target cells ^50, 51^, but so too are synthetic lipid vesicles, often referred to as lipid nanoparticles (LNPs) ^52^. To better understand the dynamics of mRNA delivery by these two natural and synthetic forms of soluble vesicles, we generated matched formulations of mRNA-loaded exosomes and mRNA-loaded LNPs. Exosomes were purified from the culture of 293F cells (***Fig. 1***), LNPs ^52^ were obtained from a commercial provider, and equal amounts of each (by vesicle number) were loaded with a synthetic mRNA encoding the hybrid luciferase/fluorescent protein Antares2 (Antares2 is comprised of the luciferase teLuc fused to two copies of the fluorescent protein CyOFP1 (CyOFP1-teLuc-CyOFP1), emits far-red shift light via bioluminescent resonance energy transfer^53^). Equal amounts of these matched exo-mRNA and LNP-mRNA formulations were then incubated at low and high doses with human cells, followed by an overnight incubation to allow for Antares2 protein expression. The next day, the cells were incubated with diphenylterazine (DTZ), a cell-permeable substrate (luciferin) for Antares2, and assayed for DTZ-dependent, Antares2-catalyzed light emission (***Fig. 2***). At low-dose administration, Antares2 expression was 25% higher in cells treated with the exo-mRNA formulation than with the LNP-mRNA formulation (n = 6, *p* = 0.0016). The difference in Antares2 expression was even more pronounced at high-dose administration, as the exo-mRNA-treated cells expressed far more Antares2 activity than the LNP-exo-treated cells (16-fold; n = 6; *p* = 0.00035).

**Figure 1.**
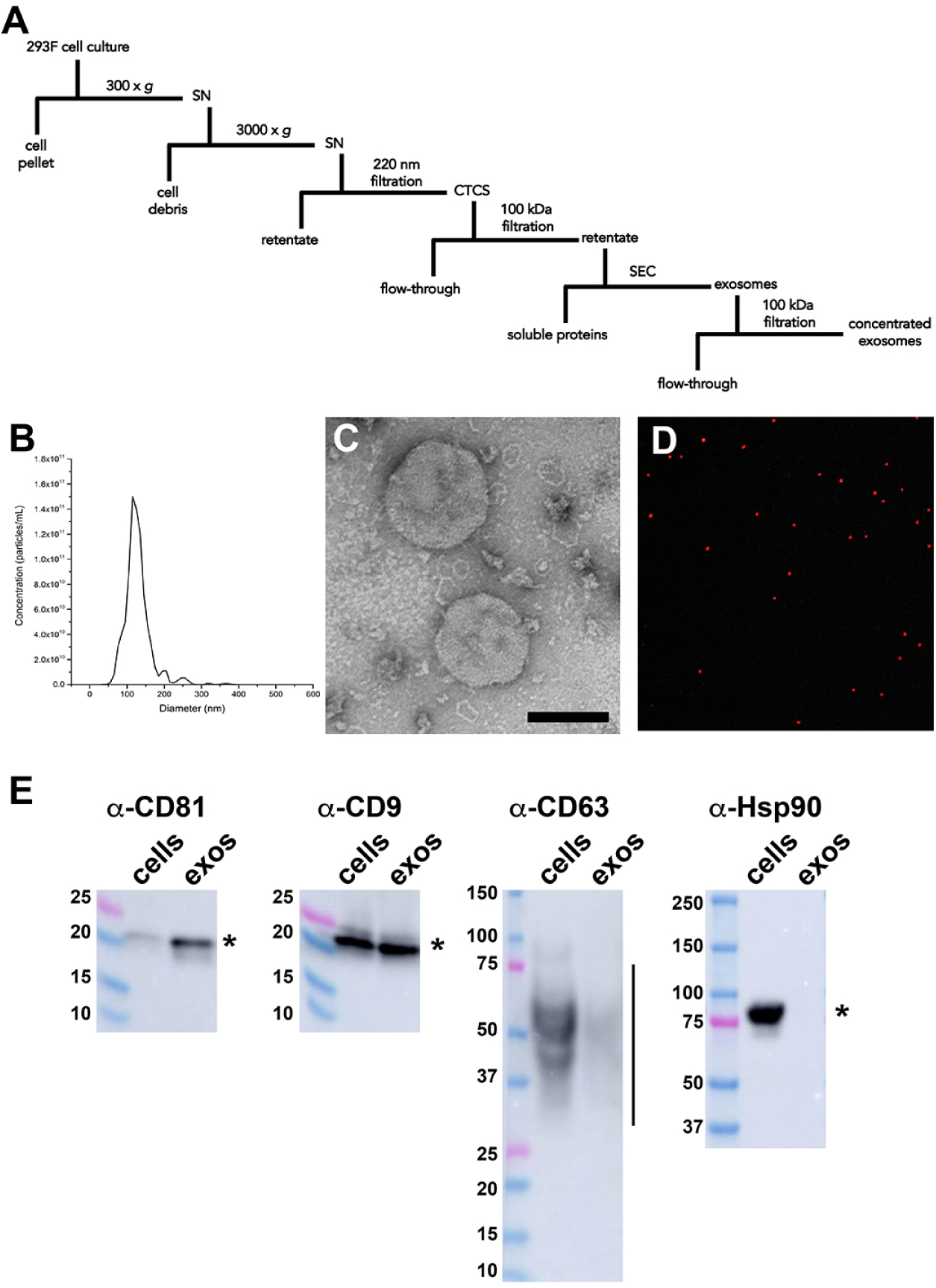
Exosome purification and characterization. (A) Schematic of exosome purification from cultures of 293F cells grown in chemically defined media. (B) NTA analysis of purified exosomes showed a mean exosome diameter of ∼115 nm. (C) Negative stain electron micrograph of purified exosomes. Bar, 100 nm. (D) Immunofluorescent NTA analysis of 293F-derived exosomes that had been labeled previously using fluorescently labeled anti-CD63 antibody. (E) Immunoblot analysis of equal proportions of 293F cell and exosome lysates using antibodies specific for the exosomal markers CD81, CD9, & CD63, as well as the control cytoplasmic protein Hsp90.

**Figure 2.**
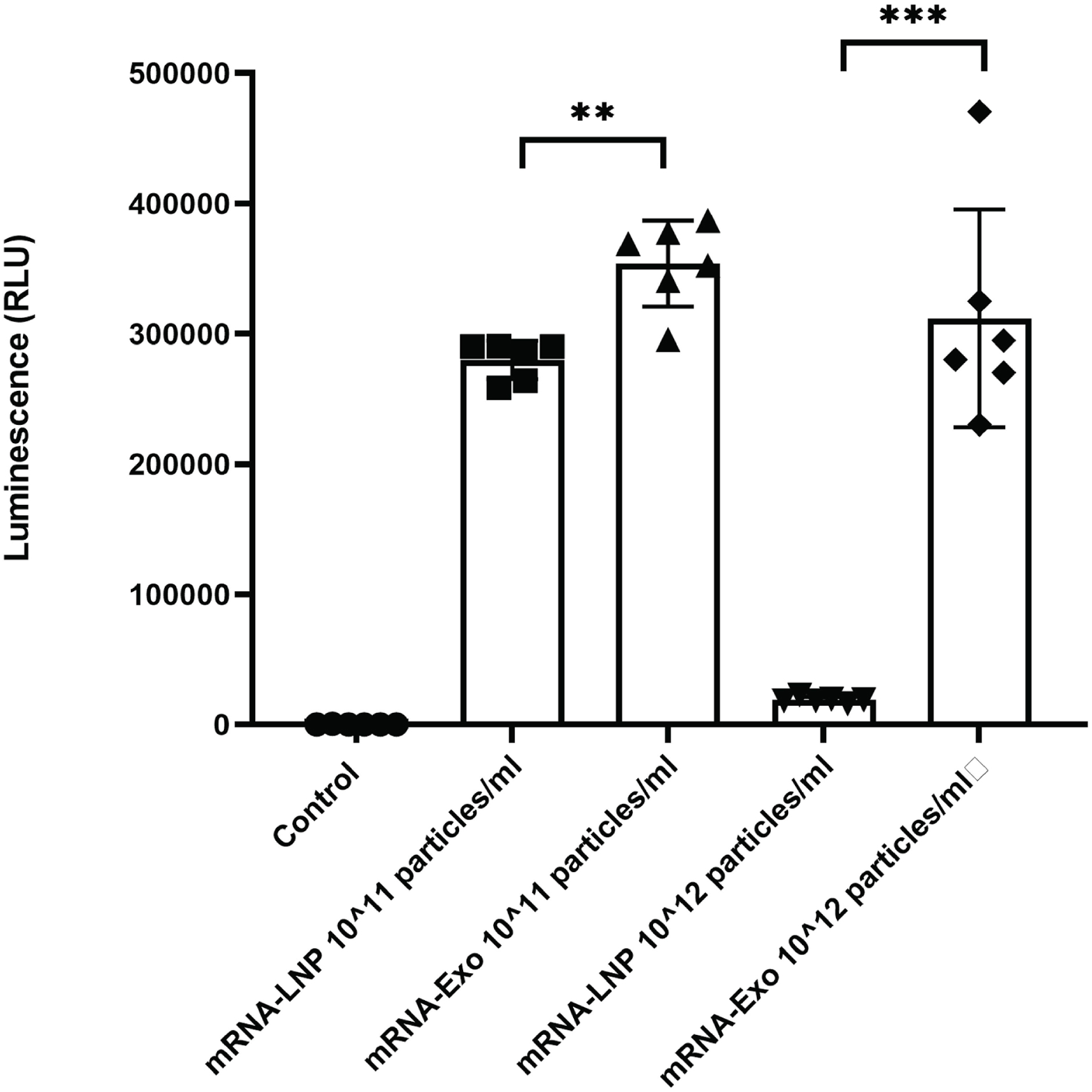
Exosomes display superior mRNA delivery characteristics. Relative luciferase activities (average +/− standard error of the mean) of cells treated with low or high concentrations of mRNA-loaded exosomes or mRNA-loaded LNPs.

This large difference in particle-mediated Antares2 expression was caused by a drop in LNP-mRNA-mediated expression, raising the possibility that LNP administration is inhibitory at high levels of administration. This in turn raised the possibility of general toxicity of LNP administration, which we addressed by following the short-term consequences of exosome and LNP injections in mice. Animals were injected (i.m.) with equal numbers of either exosomes or LNPs (50 ml ofparticles/ml), returned to their cages for three days, and then sacrificed and processed for organ histology by an independent testing laboratory (***Fig. 3A***). No abnormalities were detected in control animals (5/5) or in animals injected with exosomes (5/5). In contrast, only one of the LNP-injected animals (1/5) displayed normal spleen histology, as 4/5 animals showed an increase in red pulp. Adverse LNP effects may also explain the ∼5% reduction in body mass (n = 5; *p* = 0.05) we observed at 3 days post-injection (***Fig. 3B***).

**Figure 3.**
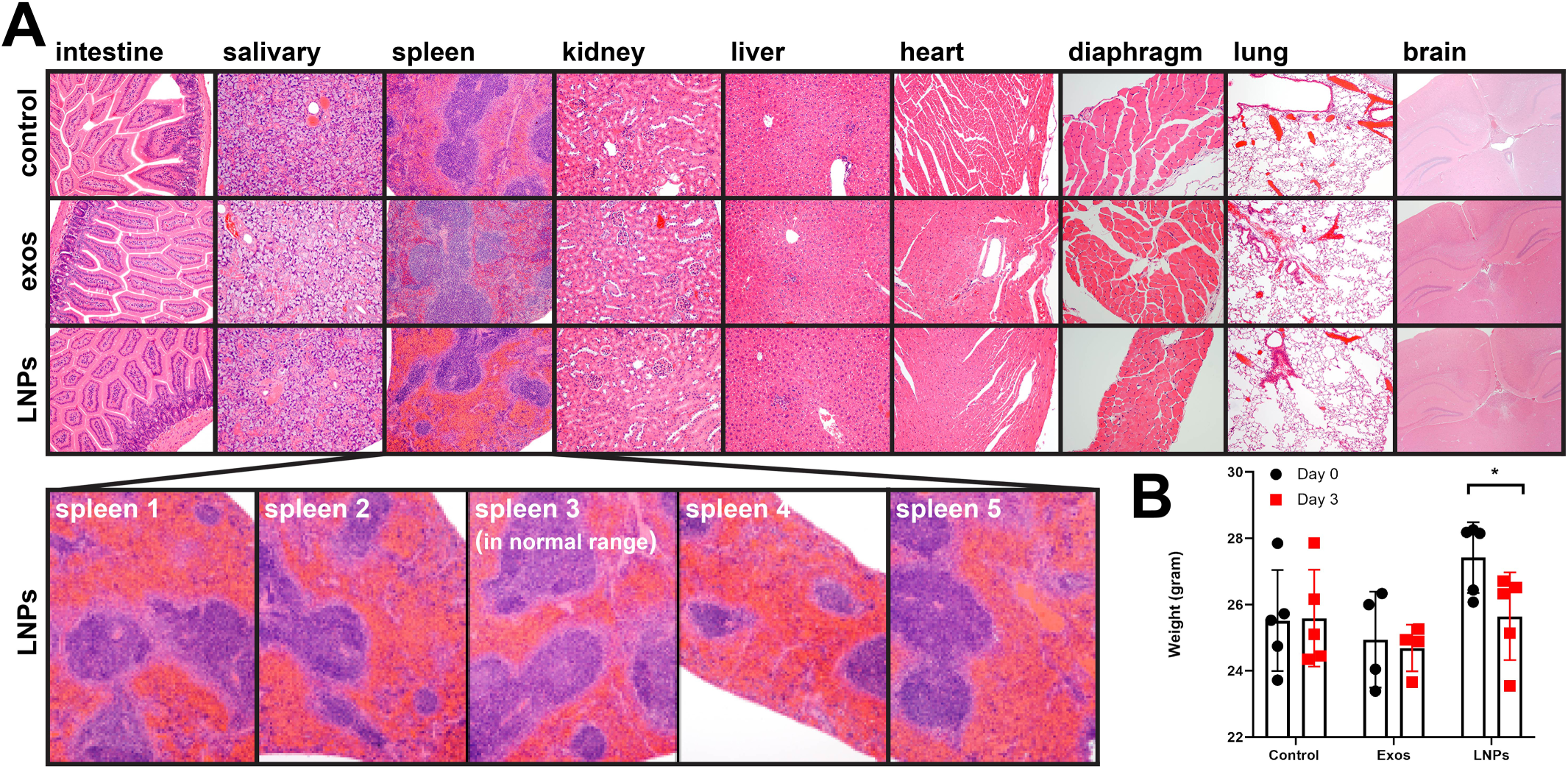
Effect of exosome and LNP injections on organ histology and body mass. (A) H&E staining of tissue sections from BALB/c mice that had been injected three days earlier with 50 ml of PBS, exosomes (10^12^/ml), or LNPs (10^12^/ml). (B) Body mass measurements prior to and at 3 days after injection. All animals were subjected to analysis by an independent pathology service, which noted spleen abnormalities in 4/5 LNP-treated animals but no abnormalities in control or exosome-treated animals.

The robust expression of exosome-delivered mRNA *in vitro* and the absence of exosome-associated adverse effects led us to next test whether RNA-loaded exosomes might also be able to drive Antares2 expression *in vivo*. Towards this end, we injected adult mice (0.05 ml volume, intramuscular (i.m.) administration) with Antares2 mRNA-loaded exosomes, returned the animals to cages to allow for Antares2 expression. 24 hours later, the control (uninjected) and treated mice were injected (i.p.) with a solution of the Antares2 luciferin DTZ and imaged immediately using a real-time bioluminescent imaging (BLI) system to visualize exosome-mediated, mRNA-directed Antares2 expression. Control animals displayed no significant light emission upon DTZ injections whereas animals that had been injected with the mRNA-loaded exosome formulation displayed robust light emission (***Fig. 3***). These observations demonstrate that RNA-loaded exosomes can deliver functional mRNAs into cells in live animals in a way that leads to mRNA translation, protein expression, and directed enzyme activity.

### Design and validation of S^W1^ and LSNME mRNAs

We next tested whether exosome-mRNA formulations can be used to elicit immune responses to mRNA-encoded antigens. Towards this end, we synthesized a pair of mRNAs, one of which expresses the form of SARS-CoV-2 Spike (S^W1^) encoded by the initial viral isolate ^1^. The second mRNA expresses a fusion protein (LSNME) comprised of the SARS-CoV-2 Nucleocapsid protein, as well as fragments of the Spike, Membrane, and Envelope proteins, all inserted in the extracellular domain of human Lamp1 (this Lamp1-based fusion protein aims to induce anti-SARS-CoV-2 immunity by targeting viral protein fragments to the MHC Class I and II antigen presentation pathways ^54, 55, 56^). Transfection of these mRNAs into HEK293 cells (***Fig. 4***) resulted in expression of Spike at the cell surface but also at internal organelles (shown elsewhere to be lysosomes ^57^), whereas expression of LSNME led to its accumulation in what appears to the endoplasmic reticulum, the site of MHC Class I peptide loading and maturation.

**Figure 4.**
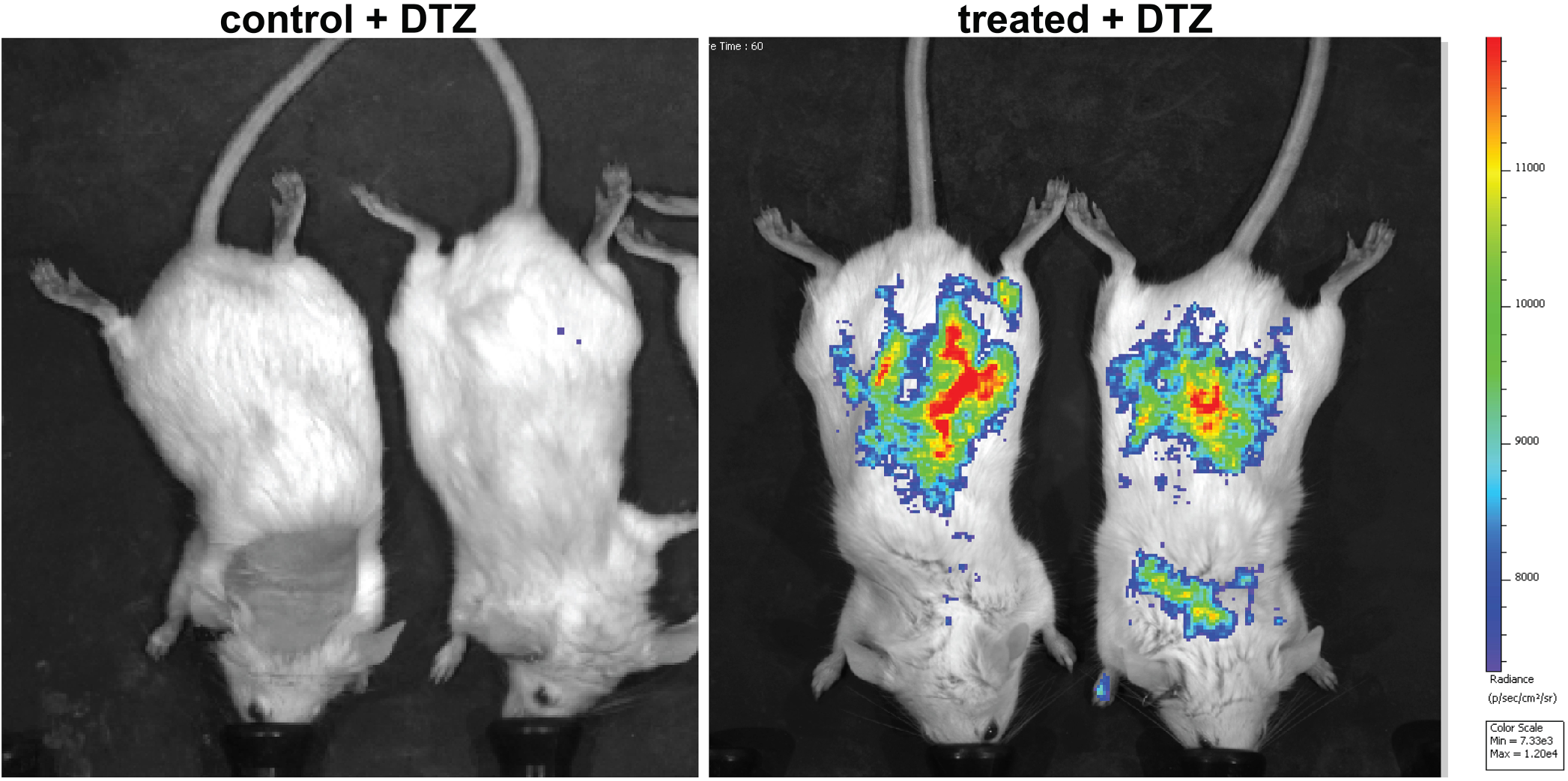
Real-time imaging of exosome-mediated, functional mRNA delivery. Combined bioluminescent and light images of control mice and treated mice immediately following i.p. administration of DTZ. Treated mice had been injected with Antares2 mRNA-loaded exosomes 24 hours prior to imaging. Radiance is in photons/second/area (cm^2^)/steradian.

### The LSNME/S^W1^ vaccine induces antibody responses to N and S

A single exosome-mRNA formulation containing both the LSNME and S^W1^ mRNAs (hereafter referred to as the LSNME/S^W1^ vaccine) was injected (i.m.) into 13 weeks-old male C57BL/6J mice (***Fig. 5***). The vaccine was dosed at 4 ug or 0.25 ug equivalents of each mRNA and injections were performed on day 1 (primary immunization), day 21 (1^st^ boost), and day 42 (2nd boost). Blood (0.1 mL) was collected on days 14, 35, 56, 70 and 84. On day 84 the animals were sacrificed to obtain tissue samples for histological analysis and splenocytes for blood cell studies. Using ELISA kits adapted for the detection of mouse antibodies, we observed that vaccinated animals displayed a dose-dependent antibody response to both the SARS-CoV-2 N protein and S protein. These antibody reactions were not particularly robust but they were long-lasting, persisting to 7 weeks after the final boost with little evidence of decline. It should be noted that the modest antibody production was expected in the case of the N protein, as the LSNME mRNA is designed to stimulate cellular immune responses rather than the production of anti-N antibodies.

**Figure 5.**
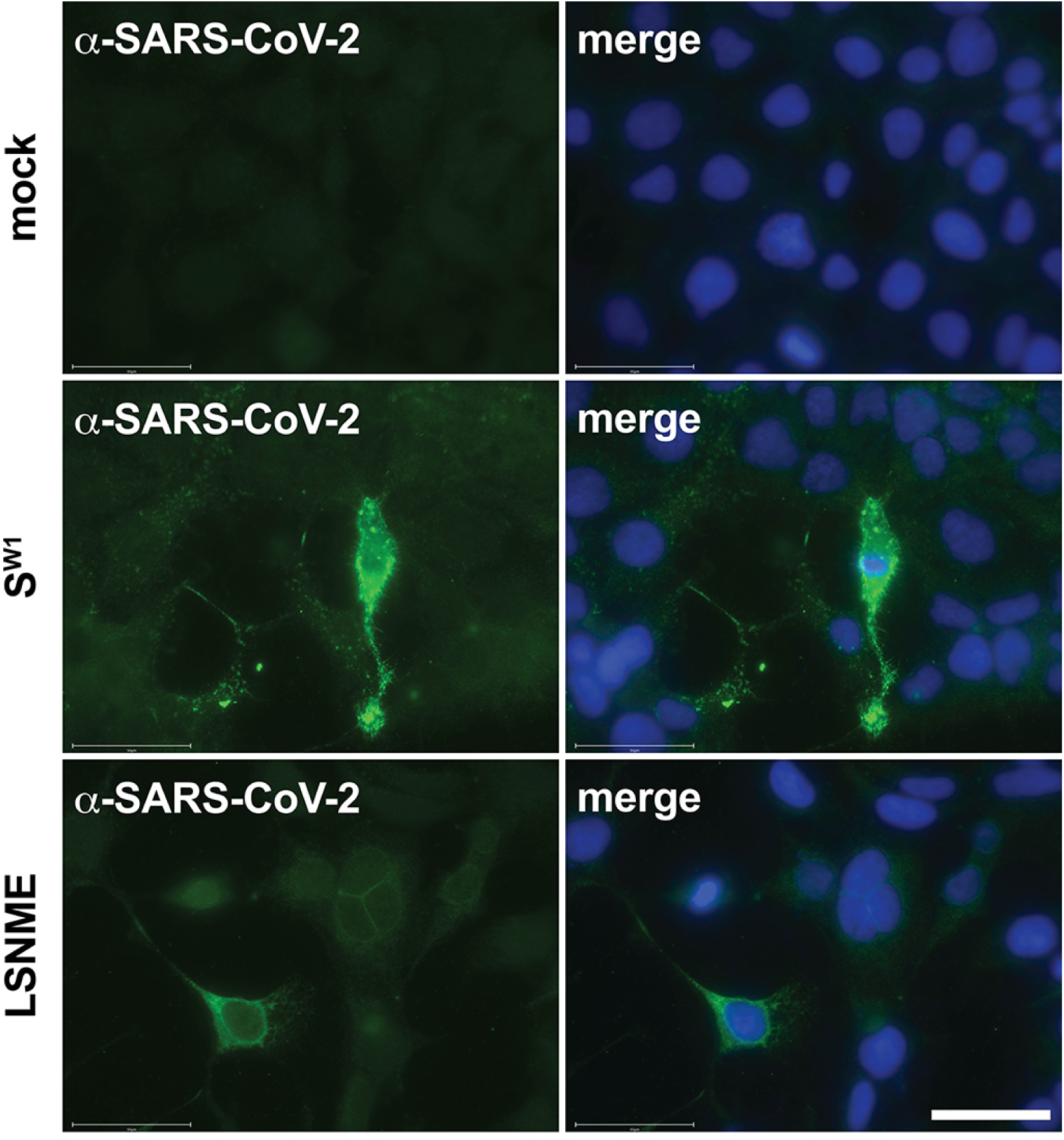
Expression of SW1 and LSNME following mRNA transfection. (A, B) Fluorescence micrographs of HEK293 cells stained with DAPI and a plasma from a COVID-19 patient. (C-F) Fluorescence micrographs of HEK293 cells stained with DAPI and plasmas from a COVID-19 patient following their transfection with the (C, D) S^W1^-encoding mRNA and (E, F) the LSNME-encoding mRNA. Bar, 50 µm.

### LSNME/S^W1^ vaccination induces cellular immune responses to N and S

Vaccinated and control animals were also interrogated for the presence of antigen-reactive CD4+ and CD8+ T-cells. This was carried out by collecting splenocytes at the completion of the trial (day 84) using a CFSE proliferation assay in the presence or absence of recombinant N and S proteins. These experiments revealed that vaccination had induced a significant increase in the percentages of CD4^+^ T-cells and CD8^+^ T-cells that proliferated in response to addition of either recombinant N protein or recombinant S protein to the culture media (***Fig. 6A-D***). These vaccine-specific, antigen-induced proliferative responses demonstrate that the LSNME/S^W1^ vaccine achieved its primary goal, which was to prime the cellular arm of the immune system to generate N-reactive CD4^+^ and CD8^+^ T-cells, and also S-reactive CD4^+^ and CD8^+^ T-cells. In additional experiments, we stained antigen-induced T-cells cells for the expression of interferon gamma (IFNγ) and interleukin 4 (IL4). These experiments revealed that the S-reactive CD4^+^ T-cell population displayed elevated expression of the Th1-associated cytokine IFNγ, and to a lesser extent, the Th2-associated cytokine IL4 (***Fig 7***). In contrast, N-reactive T-cells failed to display an N-induced expression of either IFNγ or IL4.

**Figure 6.**
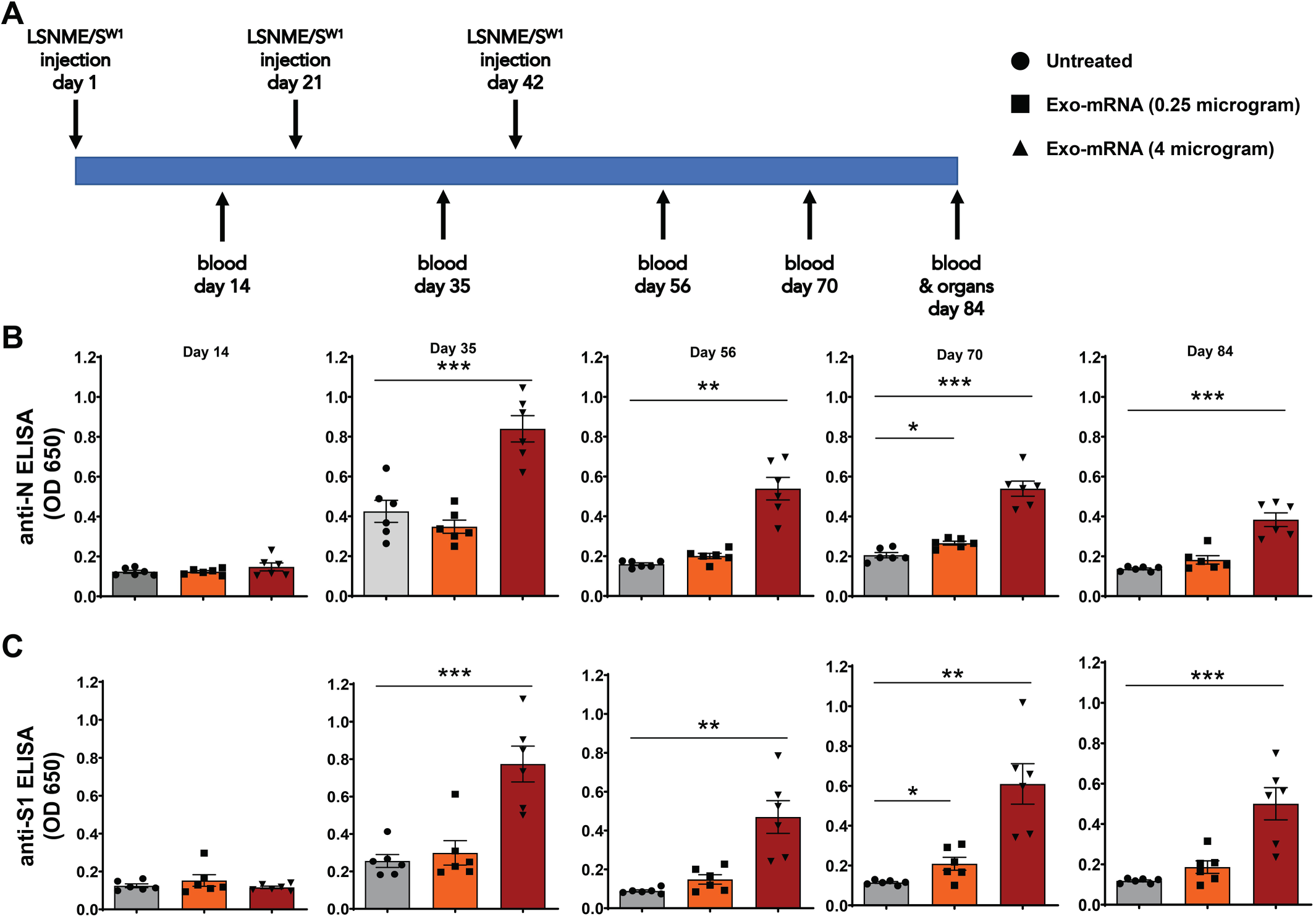
LSNME/S^W1^ vaccination induces antibody responses to SARS-CoV-2 N and S protein. (A) Schematic of immunization and blood/tissue collection timeline. (B) Anti-N ELISA results of diluted plasma from (grey bars and black circles) individual six control mice, (orange bars and black squares) six mice immunized with 0.25 μg equivalents of each mRNA, and (rust bars and black triangles) six mice immunized with 4 μg equivalents of each mRNA. (C) Anti-S1 ELISA results of diluted plasma from (grey bars and black circles) individual six control mice, (orange bars and black squares) six mice immunized with 0.25 μg equivalents of each mRNA, and (rust bars and black triangles) six mice immunized with 4 μg equivalents of each mRNA. Height of bars represents the mean, error bars represent +/− one standard error of the mean, and the statistical significance of differences between different groups is reflected in Student’s t-test values of * for <0.05, ** for <0.005, and *** for <0.0005.

**Figure 7.**
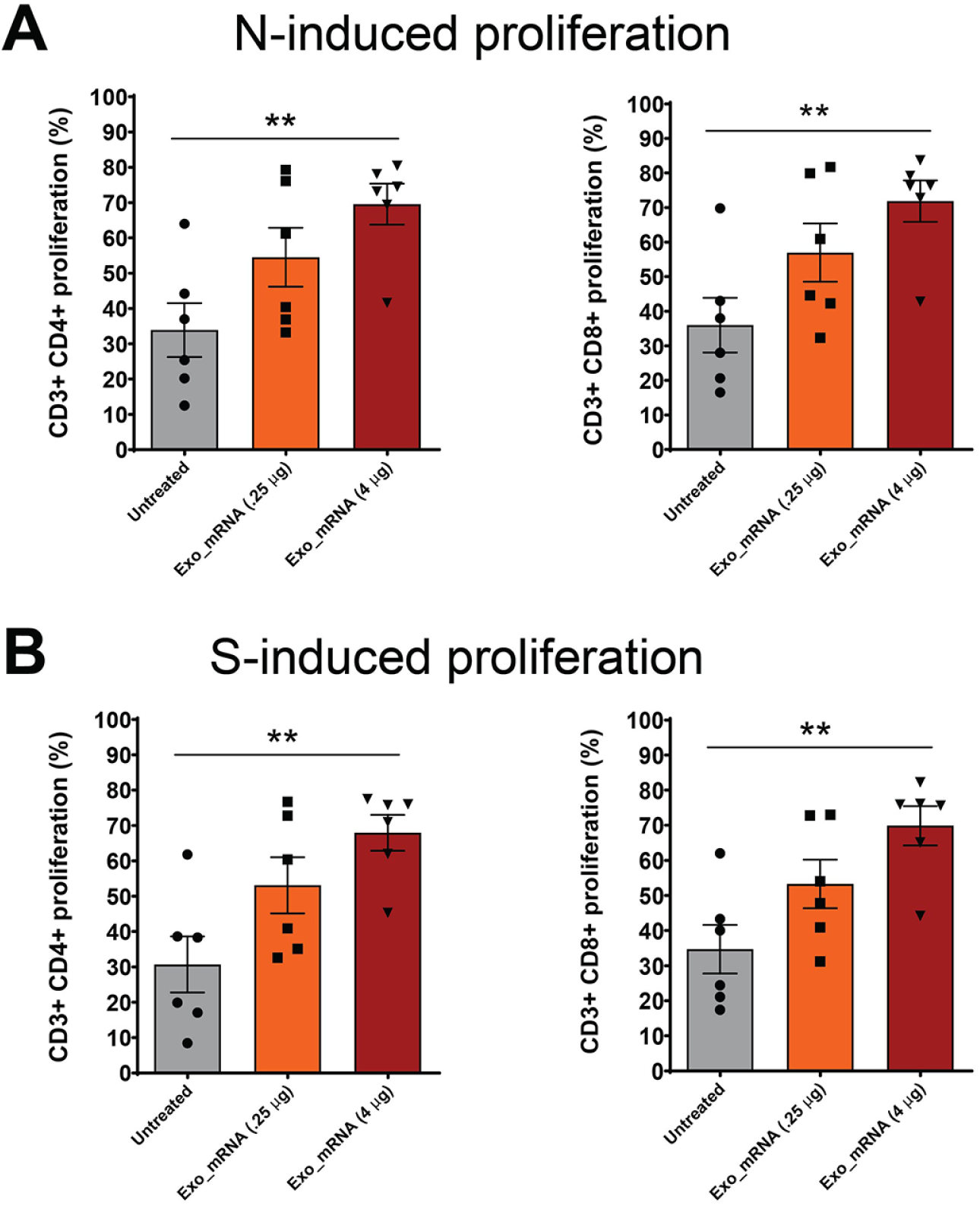
LSNME/S^W1^ vaccination induces CD4^+^ and CD8^+^ T-cell responses. CFSE-labeled splenocytes were interrogated by flow cytometry following incubation in the absence or presence of (A, B) purified, recombinant N protein or (C, D) purified, recombinant S protein, and for antibodies specific for CD4 and CD8. Differences in proliferation of CD4^+^ cells and CD8^+^ cells were plotted for (grey bars and black circles) individual six control mice, (orange bars and black squares) six mice immunized with 0.25 μg equivalents of each mRNA, and (rust bars and black triangles) six mice immunized with 4 μg equivalents of each mRNA. Height of bars represents the mean, error bars represent +/− one standard error of the mean, and the statistical significance of differences between different groups is reflected in Student’s t-test values of * for <0.05 and ** for <0.005.

### Absence of vaccine-induced adverse reactions

Control and vaccinated animals were examined regularly for overall appearance, general behavior, and injection site inflammation (redness, swelling). No vaccine-related differences were observed in any of these variables, and animals from all groups displayed similar age-related increases in body mass (***supplemental figure 1***). Vaccination also had no discernable effect on blood cell counts (***supplemental figure 2***). Histological analyses were performed on all animals at the conclusion of the study by an independent histology service, which reported that vaccinated animals showed no difference in overall appearance of any of the tissues that were examined. Representative images are presented for brain, lung, heart, liver, spleen, kidney, and side of injection skeletal muscle in an animal from each of the trial groups (***Fig. 8***).

**Figure 8.**
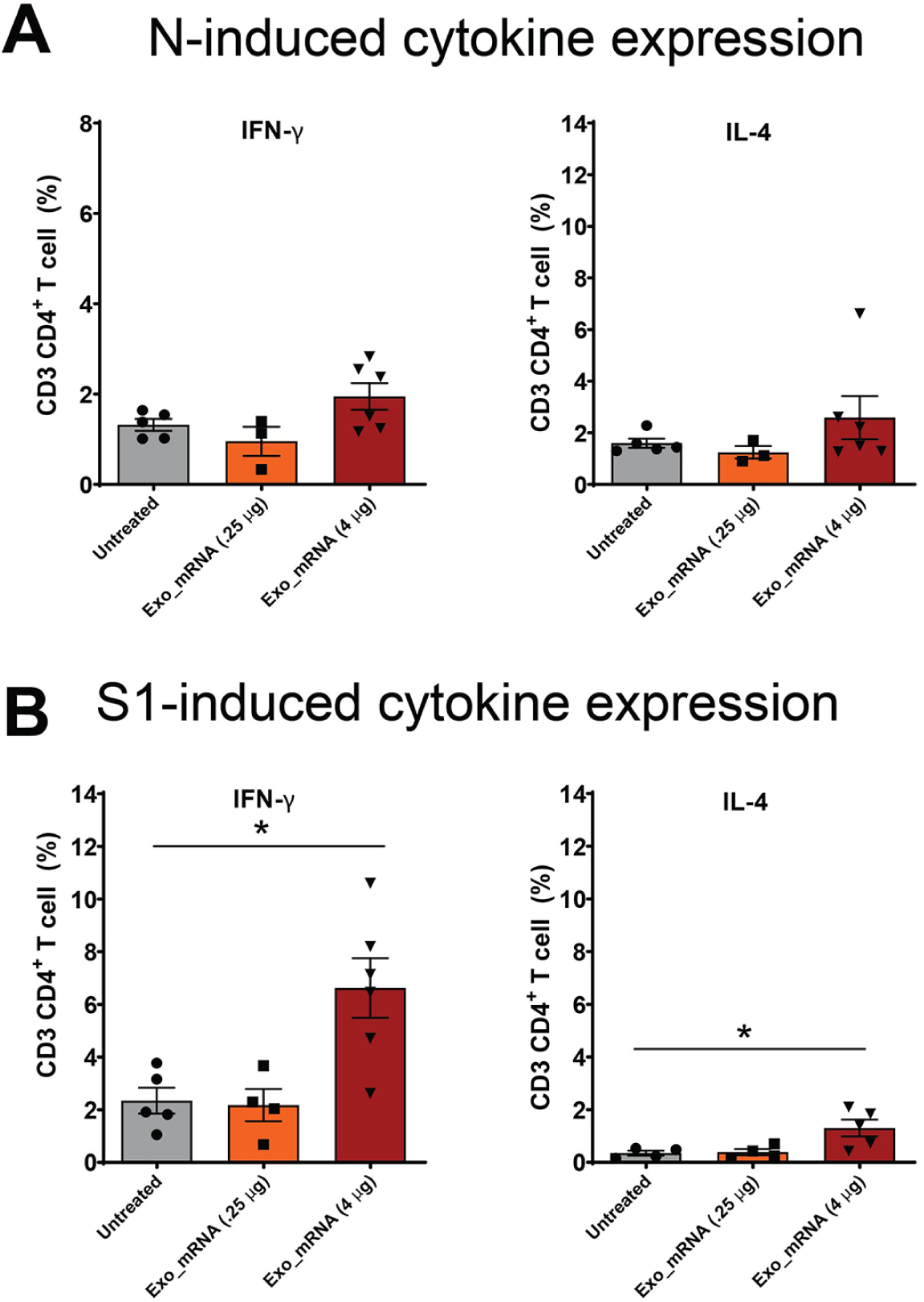
LSNME/S^W1^ vaccination leads to S-induced expression of IFNγ and IL4 by CD4^+^ T-cells. Splenocytes were interrogated by flow cytometry following incubation in the absence or presence of (A, B) purified, recombinant N protein or (C, D) purified, recombinant S protein, and labeling with antibodies specific for CD4 or CD8, and for IFNγ or IL4. Differences in labeling for IFNγ or IL4 in CD4^+^ CD8^+^ cell populations were plotted for (grey bars and black circles) individual six control mice, (orange bars and black squares) six mice immunized with 0.25 μg equivalents of each mRNA, and (rust bars and black triangles) six mice immunized with 4 μg equivalents of each mRNA. Height of bars represents the mean, error bars represent +/− one standard error of the mean, and the statistical significance of differences between different groups is reflected in Student’s t-test values of * for <0.05.

**Figure 9.**
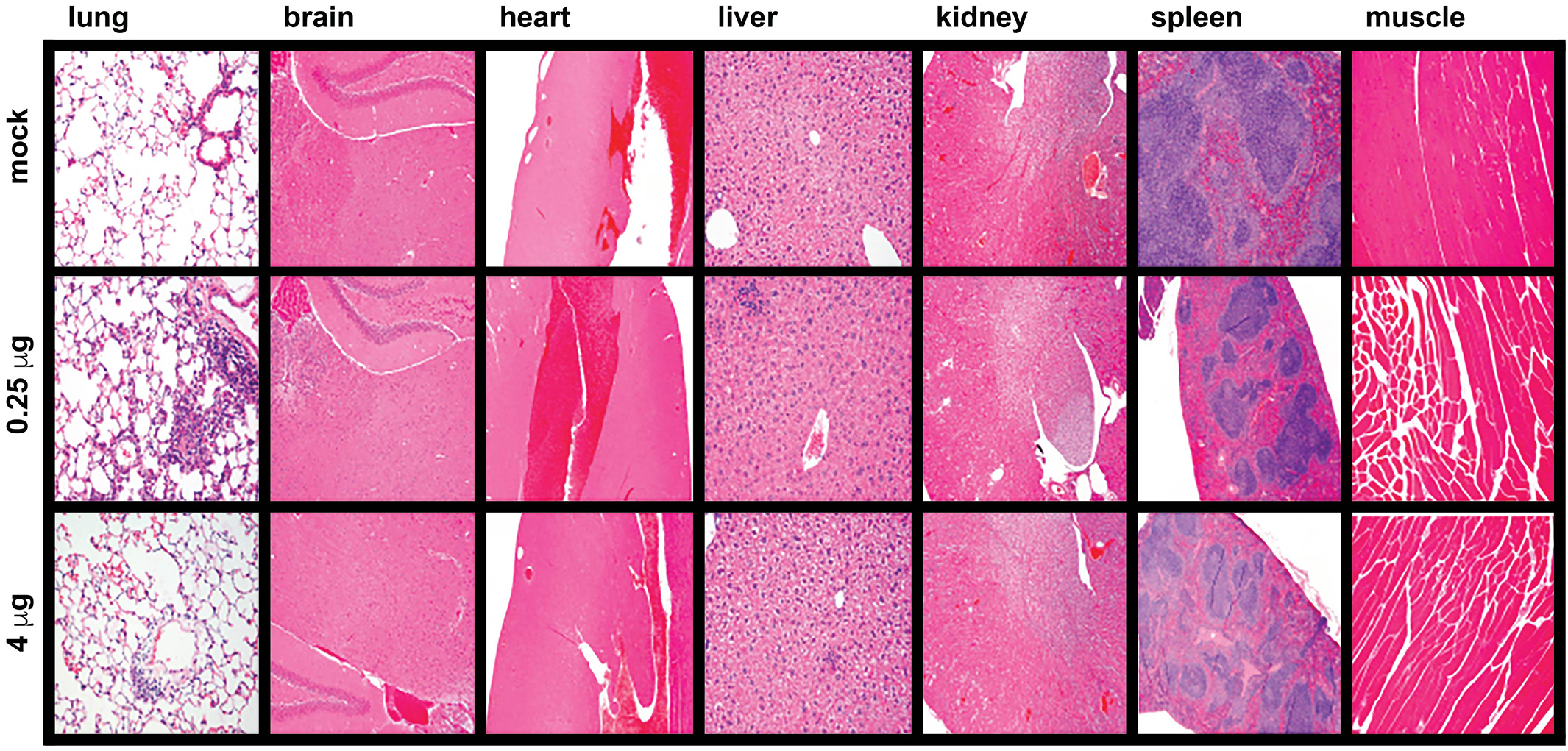
Absence of tissue pathology upon LSNME/S^W1^ vaccination. Representative micrographs from histological analysis (hematoxylin and eosin stain) of lung, brain, heart, liver, kidney, spleen, and muscle (side of injection) of animals from (upper row) control mice, (middle row) mice immunized with the lower dose of the LSNME/S^W1^ vaccine, and (lower row) mice immunized with the higher dose of the LSNME/S^W1^ vaccine.

## Discussion

Exosomes are natural products of human cells that are more ‘self’ than ‘non-self’. Immune systems are tolerant of the high levels of exosomes that are continuously present in all biofluids (e.g. blood, lymph, cerebrospinal fluid, vitreous, interstitial fluids, etc.) ^49, 58^. Furthermore, there is no evidence of adverse effects of allogeneic exosome transfer, whether of purified exosomes (from amniotic fluid, blood, etc.) or of inadvertent exosome transfer during tissue transplantation, blood transfusion, plasma injection, etc. In this context, the fact that exosomes normally participate in pathways of vesicle-mediated, intercellular RNA traffic ^59–61^ indicates that exosomes may be an ideal vehicle for clinical RNA delivery. The data presented here support this hypothesis by showing that that exosome-mRNA formulations can support the *in vivo*, functional expression of proteins as diverse as soluble cytoplasmic enzymes, viral structural proteins, and synthetic fusion proteins.

Our findings are also relevant to the ongoing battle against SARS-CoV-2. Current vaccine strategies are all centered on inducing immunity to Spike, but Spike-only vaccines are susceptible to escape effects whenever and antigenically shifted Spike variants starts to spread in susceptible populations. While we are developing strategies designed to address this challenge by improved design of expression-dependent Spike vaccines, we are also working to address it by generating a multiplexed mRNA vaccine that delivers two or more mRNAs, one encoding Spike and the others encoding Nucleocapsid and perhaps fragments of other proteins as well. One limitation of this approach is that Nucleocapsid is a cytoplasmic protein rather than a surface antigen, a topology that limits its efficacy in vaccination studies. However, this limitation can be overcome by expressing Nucleocapsid as part of a fusion with the lysosomal resident protein Lamp1, which places Nucleocapsid protein in the correct compartments for Class I and Class II antigen presentation (ER and lysosome/MHC Class II compartment, respectively). This approach was realized in our LSNME/S^W1^ vaccine, which elicited strong cellular immune responses to Nucleocapsid as well as to Spike. Vaccinated animals displayed antigen-induced CD4^+^ and CD8^+^ T-cell responses to both Nucleocapsid and to Spike that persisted for nearly two months after immunization. Furthermore, when these cell populations were interrogated for antigen-induced expression of the cytokines IFNγ and IL4, we detected elevated expression of IFNγ in CD4+ T-cells exposed to exogenous Spike protein, as well as a more modest Spike-induced expression of IL4. These results raise the possibility that the exosome-based LSNME/S^W1^ vaccine induces the kind of Th1-skewed cellular immune response desired for an anti-viral vaccine. Vaccinated animals also developed durable antibody responses to the Nucleocapsid and the Spike proteins that were sustained at relatively constant levels over the 7 weeks following immunization. This multi-antigen immune response bodes well for this approach in the next generation of SARS-CoV-2 vaccines that will be needed to protect against the emerging array of antigenically distinct SARS-CoV-2 viral strains and their ever-increasing spectrum of Spike protein mutations.

In conclusion, the results presented in this study validate the use of multiplexed exosome-mRNA formulations for functional delivery of mRNAs both in cultured cells and in live animals. The successful use of exosomes to deliver Antares2 mRNA opens the door to follow-on studies aimed at optimizing exosome-RNA formulation conditions, as well as for characterizing the time-dependence of Antares2 expression, biodistribution of exosome-mediated RNA expression, injection site effects, and exosome-mediated tissue tropism. As for the future development of exosome-based SARS-CoV-2 mRNA vaccines, we anticipate that follow-on studies will demonstrate multiple advantages of exosome-based delivery, improved antigen designed, and most importantly, improved protective effects that arise from immunization with multiple viral antigens, and particularly Nucleocapsid, which is a main target of anti-SARS-CoV-2 immunity in COVID-19 patients ^48^ and has proven effective in vaccine studies of other coronaviruses ^54^. Furthermore, the fact that exosomes can be deployed at high concentrations without adverse effects on cells or animals bodes well for their future use in dosing regimens that require higher-level or ongoing repeated injections.

## Methods

### Cell culture

293F cells (Gibco, Cat.# 51-0029) were tested for pathogens and found to be free of viral (cytomegalovirus, human immunodeficiency virus I and II, Epstein Barr virus, hepatitis B virus, and parvovirus B19), and bacterial (*Mycoplasma*) contaminants. Cells were maintained in FreeStyle 293 Expression Medium (Gibco, #12338-018) and incubated at 37°C in 8% CO_2_. For exosome production, 293F cells were seeded at a density of 1.5 x 10^6 cells/ml in shaker flasks in a volume of ∼1/4 the flask volume and grown at a shaking speed of 110 rpm. HEK293 cells were grown in Dulbecco’s modified Eagle’s medium supplemented with 10% fetal calf serum.

### Exosome purification

293F cells were grown in shaker cultures for a period of three days. Cells and large cell debris were removed by centrifugation at 300 x *g* for 5 minutes followed by 3000 x *g* for 15 minutes. The resulting supernatant was passed through a 0.22 µm sterile filtration filter unit (Thermo Fisher, #566-0020) to generate a clarified tissue culture supernatant (CTCS). The CTCS was concentrated by centrifugal filtration (Centricon Plus-70, Ultracel-PL Membrane, 100 kDa size exclusion, Millipore Sigma # UFC710008), with ∼120 mLs CTCS concentrated to ∼0.5 mLs. Concentrated CTCS was then purified by size exclusion chromatography (SEC) in 1x PBS (qEV original columns/35 nm: Izon Science, #SP5), with the exosomes present in each 0.5 mL starting sample eluting in three 0.5 mL fractions. Purified exosomes were reconcentrated using Amicon^®^ Ultra-4 100 kDa cutoff spin columns (#UFC810024). This process yielded a population of exosomes/small EVs that have the expected ultrastructure and size distribution profile of human exosomes and contain the exosomal marker proteins CD9 and CD63 (***Fig. 8***), at a concentrating effect of ∼500-fold, to a final concentration of ∼2 x 10^12^ exosomes/ml, representing an average recovery of 35%.

### Nanoparticle Tracking Analysis (NTA)

Vesicle concentrations and size distribution profiles of exosome preparations were measured by nanoparticle tracking analysis (NTA) using a NanoSight NS300 (Malvern Panalytical, United Kingdom) in 1x PBS clarified by filtration through a 0.22 µm sterile filtration unit. Measurements were carried out in triplicates at ambient temperature with fixed camera settings (level of 14, screen gain of 10, detection threshold 3, and temperature of 21.7-22.2 °C). Immunostaining nanoparticle tracking analysis (NTA) was performed using fluorescently labeled antibody conjugate directed against human CD63 (AlexaFluor488-conjugated clone 460305; R&D Systems (Minneapolis, USA)). The fluorescently labeled anti-CD63-antibody (1 μl) was incubated with exosomes (9 μl) for 2 hours at room temperature in the dark, then diluted by addition of 1 ml of sterile-filtered PBS (Thermo Fisher, USA) and examined for exosome abundance, size, and CD63 immunoreactivity using a Particle Metrix ZetaView® TWIN device. Samples were visualized in scatter mode using the 488 nm laser and standard instrument settings (sensitivity: 80, shutter: 100, min. brightness: 30; min. area: 10; max. area: 1000) in fluorescence mode with standard fluorescence settings (sensitivity: 88, shutter: 100, min. brightness: 25; min. area: 10; max. area: 1000). The resulting videos were analysed with the ZetaView® software 8.05.10 (Particle Metrix, Germany).

### Immunoblots

Exosome and cell lysates were separated by SDS-PAGE using pre-cast, 4-15% gradient gels (Bio-Rad 4561086) and transferred to PVDF membranes (ThermoFisher, #88518). Membranes were blocked, probed with antibodies directed against CD9 (clone HI9a; BioLegend), CD63 (MX-49.129.5), CD81 (555675; BD Pharmingen), or HSP90 (sc-13119; Santa Cruz Biotechnology), then washed, exposed to HRP-conjugates of goat secondary antibodies (Jackson Immunoresearch), washed, and processed for chemiluminescent imaging using HRP-activated chemiluminescence detection solution (Amersham ECL Western Blotting Detection Reagents; cat# RPN2106), and imaged using a GE Amersham Imager 600. Images were exported as JPEG files, analyzed using ImageJ software, and processed using Photoshop (Adobe).

### Electron Microscopy and light microscopy

Exosomes were fixed by addition of formaldehyde to a final concentration of 4%. Carbon-coated grids were placed on top of a drop of the exosome suspension. Next, grids were placed directly on top of a drop of 2% uranyl acetate. The resulting samples were examined with a Tecnai-12 G2 Spirit Biotwin transmission electron microscope (John Hopkins University, USA). Fluorescence micrographs of Antares2 expression in transfected HEK293 cells were captured as PNG files using an EVOS M7000 microscope equipped with an Olympus UPlanSAPo 40x/0.95 objective.

### Production of mRNA-loaded exosomes and LNPs

mRNAs were obtained from a commercial provider (Trilink). mRNAs were purified using RNeasy columns (Qiagen) and resuspended in DNase-free, RNase-free water using nuclease-free tips and tubes. Purified mRNAs were pre-incubated with a coating of polycationic lipids and then mixed with equal amounts of either purified exosomes or LNPs (DOTAP/DOPE, #F50102, FormuMAx Scientific Inc) at 4°C for 10 minutes. Formulations were either used immediately or frozen at −80°C and thawed rapidly prior to use.

### Luciferase measurements and bioluminescent imaging

HEK293 cells were incubated with exosome-mRNA formulations overnight under standard culture conditions. Antares2 luciferase activity was measured by Live cell bioluminescence was collected after incubating with substrate diphenylterazine (MCE, HY-111382) at final concentration of 50 µM for 3 minutes. Readings were collected using a SpectraMax i3x (Molecular Devices). For in vivo studies, thirteen months-old, female Balb/c mice (Jackson Laboratory) housed under pathogen-free conditions at the Cedars-Sinai Medical Center animal facility were used to study the expression of Exosome-Anteres2 mRNA expression 24 hours after injection. Intramuscular injections were at a volume of 50 μls per mouse containing 5 ug mRNA. After 24 hours the animals were imaged using an IVIS Spectrum imager (PerkinElmer, Waltham, MA) (All animal experimentation was performed following institutional guidelines for animal care and were approved by the Cedars-Sinai Medical Center IACUC (#8602).

### Animal experimentation

All animal experimentation was performed following institutional guidelines for animal care and were approved by the Cedars-Sinai Medical Center IACUC (#8602). All injections were at a volume of 50 μls. Experiments involved injection of exosomes, LNPS, and Antares2 mRNA-loaded exosomes were performed with BALB/c mice (Jackson Laboratory). Immunization with mRNA-loaded exosomes were performed on thirteen weeks-old, male C57BL/6J mice (Jackson Laboratory) housed under pathogen-free conditions at the Cedars-Sinai Medical Center animal facility. Blood (∼0.1 mL) was collected periodically from the orbital vein. At day 84, mice were deeply anesthetized using isoflurane, euthanized by cervical dislocation, and processed using standard surgical procedures to obtain spleen, lung, brain, heart, liver, kidney, muscle, and other tissues. Spleens were processed for splenocyte analysis, and all tissues were processed for histological analysis by fixation in 10% neutral buffered formalin. Histological analysis was performed by the service arm of the HIC/Comparative Pathology Program of the University of Washington.

### ELISA for SARS-CoV-2 antigen-specific antibody responses

Mouse IgG antibody production against SARS-CoV-2 antigens was measured by enzyme-linked immunosorbent assays (ELISA). For antigens S1 (RBD) and N, pre-coated ELISA plates from RayBiotech were utilized (IEQ-CoV S RBD-IgG; IEQ-CoVN-IgG), and the experiments were performed according to the manufacturer’s instructions, with modification. Briefly, mouse plasmas at dilutions of 1:50 were added to antigen pre-coated wells in duplicates and incubated at room temperature (RT) for 2 hours on a shaker (200 rpm). The plates were washed 4 times with wash buffer followed by blocking for 2 hours at RT with 1% BSA in PBS. Mouse antibodies bound to the antigens coated on the ELISA plates were detected using HRP-conjugated goat anti-mouse secondary antibodies (Jackson Immuno Research Inc.) Plates were washed 4 times with washing buffer, and developed using TMB substrate (RayBiotech). Microplate Reader was used to measure the absorbance at 650 nm (SpectraMaxID3, Molecular Devices, with SoftMax Pro7 software).

### Single cell splenocyte preparation

After terminal blood collection, mice were euthanized, and part of fresh spleens were harvested. Single cell splenocyte preparation was obtained by machinal passage through a 40 µm nylon cell strainer (BD Falcon, #352340). Erythrocytes were depleted using Ammonium-Chloride-Potassium (ACK) lysis buffer (Gibco, #A10492-01), and splenocytes were washed using R10 media by centrifuging at 300x *g* for 5 minutes at RT. R10 media (RPMI 1640 media (ATCC, Cat#302001) supplemented with 10% fetal bovine serum (FBS) (Atlas, #E01C17A1), 50 µM 2-mercaptoethanol (Gibco, #21985-023), penicillin/streptomycin (VWR life sciences, #K952), and 10 mM HEPES (Gibco, #15630-080)) was used for all analyses of blood cells. The cells were resuspended in fresh media and counted in hemocytometer counting chamber to be used in subsequent experiments.

### Spleen lymphocyte population characterization

Splenocytes (2 x 10^5^ cells/mouse) were resuspended in 100 µL of 10% FBS in 1x PBS and incubated with fluorochrome-conjugated antibodies for surface staining of CD3 (Invitrogen, #17-0032-82) CD4 (Biolegend, #100433), CD8 (Biolegend, #100708), B220 (BD, #552771) CD11c (Invitrogen, #17-0114-81), F4/80 (Invitrogen, #MF48004) Ly6G (Invitrogen, #11-9668-80) and Ly6C (BD, #560592)) for 30 minutes at 4 ^°^C in the dark. Following incubation, samples were washed twice with 200 µLs 10% FBS in 1x PBS and centrifuged at 300 x *g* for 5 minutes at RT to remove unbound antibodies. Next the cells were fixed with 100 µLs ICS fixation buffer (Invitrogen, #00-8222-49). Samples were analyzed on a FACS Canto II (BD Biosciences) with 2,000 – 10,000 recorded lymphocytes. The data analysis was performed using FlowJo 10 software (FlowJo, LLC) and presented as a percentage change in the immune cell population compared to the vehicle-treated group.

### SARS-CoV-2 antigen-specific T cell proliferation assay using CFSE

Splenocytes were resuspended at 10^6^ cells/mL in 10% FBS in 1xPBS and stained with carboxyfluorescein succinimidyl ester (CFSE) (Invitrogen, #C34554) by rapidly mixing equal volume of cell suspension with 10 µM CFSE in 10% FBS in 1x PBS for 5 minutes at 37°C. The labeled cells were washed three times with R10 complete medium. The cells were incubated for 96 hours in the presence of 10 µg/mL SARS-CoV-2 antigens N or S1 (Acro Biosystems, #NUN-C5227; SIN-C52H4) or medium alone as negative control. After 96 hours, cells were washed with 200 µLs 10% FBS in 1xPBS and centrifuged at 300 x *g* for 5 minutes at RT. Cells were then stained with anti-CD3-APC (Invitrogen, #17-0032-82), anti-CD4-PerCP-Cy5.5 (Biolegend, #100433), and anti-CD8-PE antibodies (Biolegend, #MCD0801) for 30 minutes at 4^°^C. The stained cells were washed twice with 200 µLs 1x PBS and analyzed on a FACS Canto II (BD Biosciences). For analysis, lymphocytes were first gated for CD3+ T-cells, then for CD4+/CD8− or CD8+/CD4− populations. The data analysis was performed using FlowJo 10 software (FlowJo LLC).

### Intracellular staining for cytokines

2.0 x 10^5^ splenocytes/mouse were incubated for 72 hours in the presence of 10 µg/mL SARs-CoV2 antigens N or S1 (Acro Biosystems) or R10 medium alone (negative control). After 72 hours, the cells were washed with fresh R10 medium and incubated with phorbol myristate acetate (PMA) at concentration of 50 ng/mL (Sigma, #P1585), ionomycin at concentration of 350 ng/mL (Invitrogen, #124222), and GogiPlug at concentration of 0.8 μL/mL (Invitrogen, #51-2301KZ) for 4 hours to amplify cytokine expression in T cells. The cells were then washed with 10% FBS in 1x PBS and stained with anti-CD3-APC, anti-CD4-PerCP-Cy5.5, and anti-CD8-PE antibodies (Added above) for 30 minutes at 4^°^C in dark. The cells were washed twice with 1xPBS followed by permeabilization step using ready-to-use buffer (Invitrogen #00-8333-56). Next the cells were fixed with ICS fixation bufferAdded above for 10 minutes at RT in dark and stained intracellular for IFN-γ (eBioscience, #11-7311-82), IL-10 (eBioscience, #11-7101-82), IL-4 (Invitrogen, #12-7041-41) and Foxp3 (Invitrogen, #12-5773-80) overnight at 4^°^C in permeabilization buffer. The stained cells were analyzed on a BD FACS Canto II with 5,000 – 10,000 recorded lymphocytes. The data analysis was performed using FlowJo 10 software.

### Statistical Analysis

Statistical analysis was performed using GraphPad Prism 8 software for Windows/Mac (GraphPad Software, La Jolla California USA) or Excel. Results are reported as mean ± standard deviation or mean ± standard error, and the differences were analyzed using Student’s t-test or one-way analysis of variance.

## Acknowledgments

We thank Omid Sheikh for outstanding technical assistance during the course of these studies.

---

**Supplemental Figure 1.**
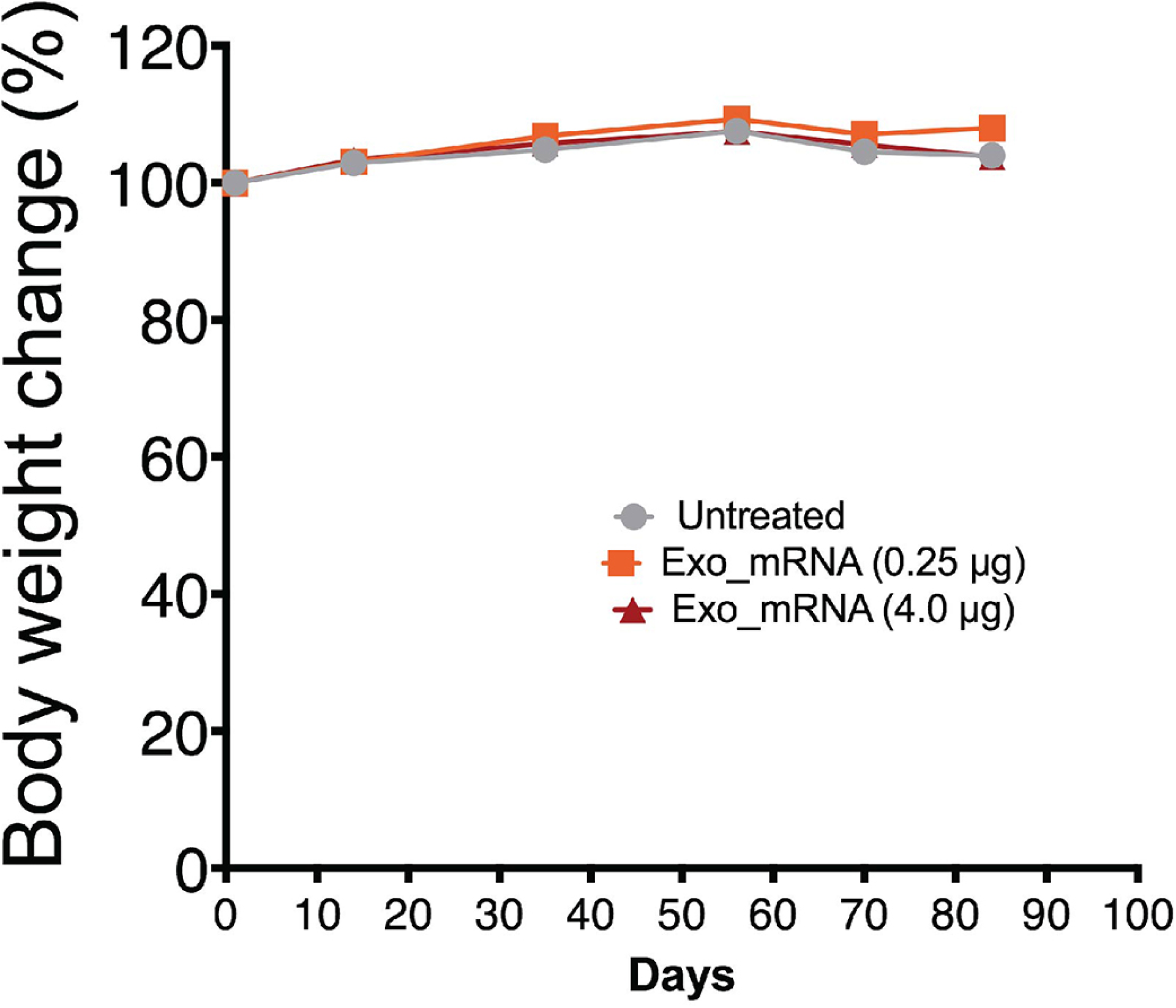
Equivalent growth of vaccinated and control animals. Body mass of all mice was measured over the course of the study and plotted as average +/− the standard error of the mean, relative to the body mass at the initiation of the trial, with groups reported as (grey lines and circles) control mice, (orange lines and squares) lower dose-treated mice, and (rust lines and triangles) higher dose-treated mice.

**Supplemental Figure 2.**
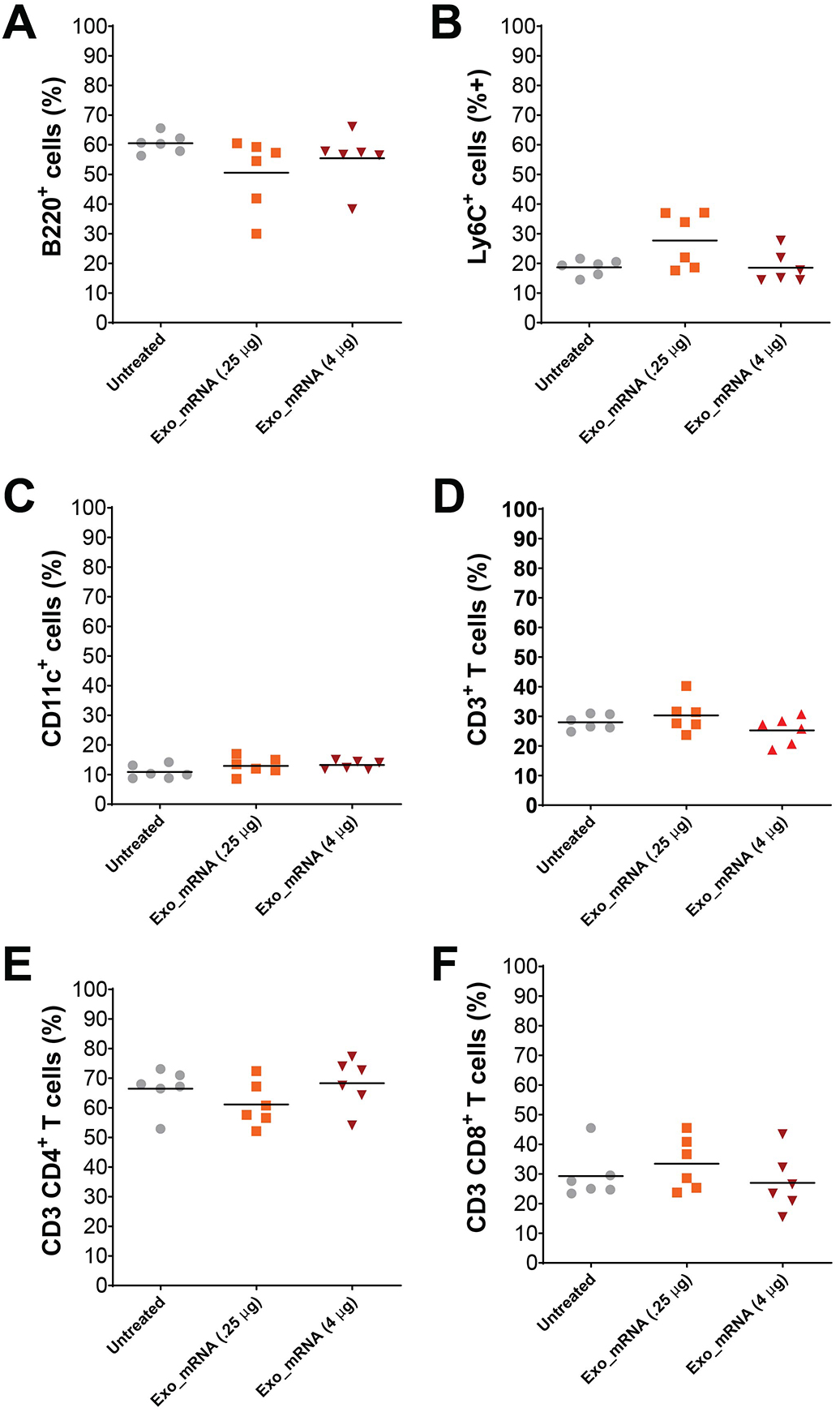
Vaccination does not induce changes in the proportional representation of key blood cell populations. Splenocytes were interrogated by flow cytometry using antibodies specific for (A) B220, (B) Ly6C, (C) CD11c, and (D) CD3. CD3^+^ cells were further differentiated by staining for (E) CD4 and (F) CD8. No statistically significant differences were detected in these subpopulations of white blood cells.

